# Alpha oscillations shape sensory representation and perceptual sensitivity

**DOI:** 10.1101/2021.02.02.429418

**Authors:** Ying Joey Zhou, Luca Iemi, Jan-Mathijs Schoffelen, Floris P. de Lange, Saskia Haegens

**Author notes:** **Corresponding author:** Saskia Haegens.

## Abstract

Alpha activity (8–14 Hz) is the dominant rhythm in the awake brain, and thought to play an important role in setting the brain’s internal state. Previous work has associated states of decreased alpha power with enhanced neural excitability. However, evidence is mixed on whether and how such excitability enhancement modulates sensory signals of interest versus noise differently, and what, if any, the consequences are for subsequent perception. Here, human subjects (male and female) performed a visual detection task in which we manipulated their decision criteria in a block-wise manner. While our manipulation led to substantial criterion shifts, these shifts were not reflected in pre-stimulus alpha-band changes. Rather, lower pre-stimulus alpha power in occipital-parietal areas improved perceptual sensitivity and enhanced information content decodable from neural activity patterns. Additionally, oscillatory alpha phase immediately before stimulus presentation modulated accuracy. Together, our results suggest that alpha-band dynamics modulate sensory signals of interest more strongly than noise.

**Significance statement:** The internal state of our brain fluctuates, giving rise to variability in perception and action. Neural oscillations, most prominently in the alpha-band, have been suggested to play a role in setting this internal state. Here, we show that ongoing alpha-band activity in occipital-parietal regions predicts the quality of visual information decodable in neural activity patterns, and subsequently human observer’s sensitivity in a visual detection task. Our results provide comprehensive evidence that visual representation is modulated by ongoing alpha-band activity, and advance our understanding on how, when faced with unchanging external stimuli, internal neural fluctuations influence perception and behavior.

## INTRODUCTION

Identical stimuli may lead to variable percepts: when presented with the same threshold-level stimulus repeatedly, an observer sometimes detects it while sometimes misses it. The brain’s internal state, largely determined by moment-to-moment neural activity fluctuations, contributes to this perceptual variability (for reviews, see Harris & Thiele, 2011; Samaha et al., 2020).

Alpha activity (8–14 Hz) is the dominant rhythm in the awake brain. One prevailing hypothesis is that the alpha rhythm sets the internal state’s excitability level via functional inhibition (Jensen & Mazaheri, 2010; Klimesch et al., 2007). Because of its tight link with internal excitability, alpha-band oscillatory activity (and its temporal fluctuations) is a plausible key source of perceptual variability. Indeed, extensive work reported that pre-stimulus alpha-band power modulates perception of simple sensory stimulations (Barne et al., 2020; Haegens et al., 2011; Hanslmayr et al., 2007; Thut et al., 2006; van Dijk et al., 2008; van Ede et al., 2011).

Only recently researchers started to investigate *how* internal excitability changes indexed by alpha oscillations manifest in subjects’ behavioral outcomes (Iemi et al., 2017; Samaha et al., 2017), using Signal Detection Theory (Macmillan & Creelman 2004). In this theoretical framework, two metrics are used to quantify the perceptual process: sensitivity (*d’*) – a metric of how well the observer distinguishes sensory signals of interest from noise, and criterion (*c*) – a metric of observer’s tendency for one perceptual decision over the others. Depending on potentially differential impacts on sensory signals of interest versus noise, internal excitability changes indexed by alpha activity may modulate different aspects of the perceptual process.

One hypothesis is that decreased alpha activity amplifies the sensory signal of interest and noise to the same extent (e.g., the gain is *additive* to the response amplitude) without making them more distinct from each other, and therefore only results in a criterion shift in the observer’s behavior (Figure 1). This view is mainly supported by empirical work correlating ongoing, as opposed to experimentally-induced, alpha activity changes with perceptual reports. Several studies showed that when pre-stimulus alpha activity was low, subjects reported higher subjective visibility (Benwell et al., 2017) and confidence (Samaha et al., 2017), and had a higher tendency to see a target (i.e., more liberal criterion), but displayed no change in perceptual sensitivity (Iemi et al., 2017; Limbach & Corballis, 2016). If a low state of alpha activity indeed amplifies signal and noise similarly and thereby shifts the observer’s criterion, a key question is whether (top-down) changes in criterion are implemented via (and/or reflected by) shifts in the alpha state.

**Figure 1.**
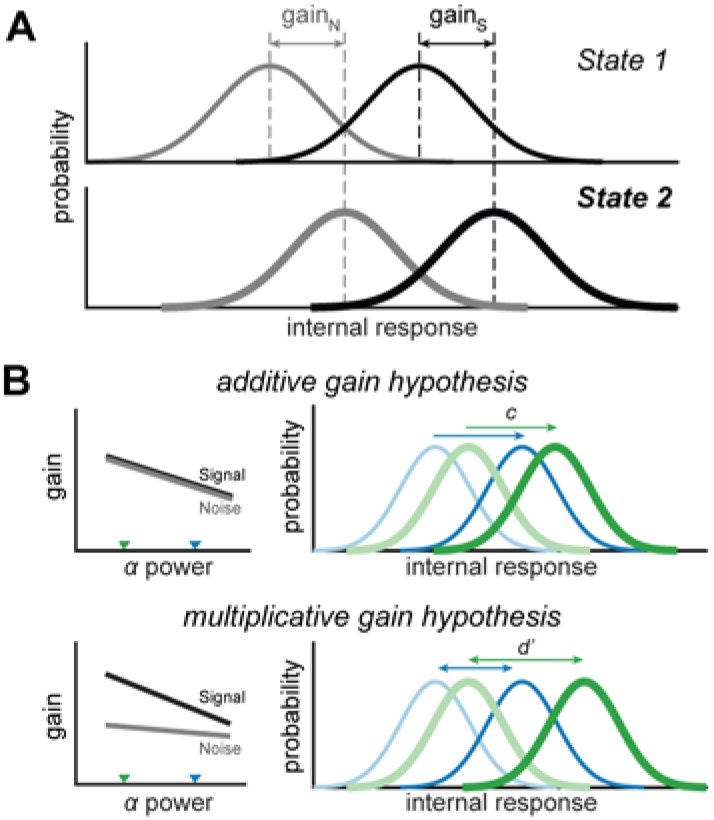
Hypothesis schematic. **(A)** Internal response distribution in different states, with state 1 the initial or baseline state (internal response distribution of signal modeled as a normal distribution with mean *μ*_*signal*1_ and standard deviation *σ*_1_, and that of noise as *μ*_*noise*1_ and *σ*_1_). We defined gain as 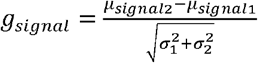 for the signal distributions, and 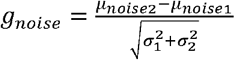 for the noise distributions. We set *σ*_1_ = *σ*_2_ here for simplicity. **(B)** Both models assume that response gain decreases as alpha power increases. The additive hypothesis predicts that the difference in response gain between signal and noise distributions remains constant across different alpha power levels, and therefore predicts a criterion shift when comparing two states of different alpha power (denoted by the triangles in the left panel). In contrast, the multiplicative hypothesis predicts that the difference in response gain scales as a function of alpha power, and therefore predicts a *d’* change when comparing two states of different alpha power.

Another hypothesis is that decreased alpha activity amplifies the signal of interest to a larger extent relative to the noise (e.g., the gain scales *multiplicatively* as a function of response amplitude), making them more distinct from each other and hence resulting in improved perceptual sensitivity (Figure 1). This view is largely inferred based on studies using classical attentional-cueing paradigms, in which pre-stimulus alpha-band activity patterns (i.e., alpha lateralization, activity difference between task-relevant and task-irrelevant regions) and perceptual sensitivity co-vary following attentional cueing (Haegens et al., 2011; Thut et al., 2006; van Ede et al., 2011). Although informative, it is unclear whether the observed correlation between alpha lateralization and performance is mainly driven by an alpha amplitude decrease in task-relevant or increase in task-irrelevant regions. Furthermore, if alpha oscillations indeed modulate sensory signal of interest more strongly relative to the noise, an immediate prediction is that decreased alpha activity leads to enhanced information coding of the task-relevant stimuli.

To differentiate between these hypotheses, we tested human subjects in a magnetoencephalography (MEG) experiment, during which we experimentally manipulated their decision criterion using a novel visual detection paradigm. We examined whether and how changes in pre-stimulus alpha dynamics (*i*) correlate with criterion shifts, (*ii*) influence sensitivity while criterion is controlled, and (*iii*) modulate information coding, indexed by the representational content decodable in the neural signal.

## MATERIALS AND METHODS

### Data availability

All data and code for stimulus presentation and analysis are available online at the Donders Repository at https://doi.org/10.34973/w1k5-sm41.

### Subjects

Thirty-four healthy volunteers completed the full experiment. Two of them were excluded from analysis because of excessive head movement during the MEG recording, resulting in the planned sample size of 32 subjects in the reported analysis (mean age = 25.8, SD = 6.0; 23 females). The study was approved by the local ethics committee (CMO Arnhem-Nijmegen). All subjects gave informed consent prior to the experiment and received monetary compensation for their participation.

### Procedure

Subjects reported to the lab on two days within one week, for a training session on day 1 and MEG recording session on day 2.

#### Training session on day 1

The training session on day 1 served to familiarize subjects with the task and to prepare them for the MEG recording session on day 2. We introduced the task with 10 “slow” trials, in which we set the ISI between the target and the backward mask to 100 ms and provided feedback at the end of each trial. Before starting the up-down staircase, subjects practiced 20 “normal-speed” trials with feedback, to get familiar with the task. This introduction phase was repeated if necessary. Two runs of 3-down-1-up staircases were then used to titrate the contrast of the backward mask, one run for each target orientation (clockwise and counterclockwise gratings; order counter-balanced across subjects). Subjects were explicitly informed about the orientation and grating presence rate (50%) at the start of each staircase run. No feedback was provided during the staircase. After the staircase procedure, subjects completed four experimental blocks. Notably, to maximize the priming effect (for the MEG session), gratings were presented 80% (20%) of the time in both the priming and main task phase, and feedback was provided at the end of each block (i.e., accuracy and mean reaction time). We aimed to convince subjects that utilizing the provided target-presence rate in their decisions was beneficial for task performance.

#### MEG testing session on day 2

On day 2, similar to the training session, subjects first completed two runs of 3-down-1-up staircases inside the MEG, to determine the contrast level of the backward mask that resulted in ~80% detection accuracy when an equal number of target-present and absent trials was presented. The staircases were done after a brief recap of the task, in which subjects completed 10 “slow” trials and 20 “normal-speed” trials with trial-by-trial feedback (i.e., same as the introduction phase of the training session on day 1). They then performed eight main experimental blocks and three grating localizer blocks, during which neurophysiological data were recorded.

### Experimental paradigm

#### Main experimental block

Two stimuli, a target and a mask, were presented sequentially in each trial, and the subject’s task was to report as accurately as possible whether a grating was present or not in the trial (see Figure 2A for trial schematic). To introduce criterion shift in the subject’s perceptual decisions, we designed a paradigm inspired by Crapse and colleagues (Crapse et al., 2018). Each main experimental block started with an instruction screen, informing subjects of the orientation (CW or CCW) and grating presence rate (20% or 80%) in the upcoming block. Every block consisted of 40 priming trials and 80 main task trials. For the first 40 priming trials, gratings were presented either 20% or 80% of the time, consistent with the block instruction. The backward masks were of 5% Michelson contrast, which had little masking effect on the target grating. We purposely presented the targets in a clearly discriminable way during the priming phase, to encourage subjects to shift their decision criteria accordingly. Before starting the 80 main task trials, a reminder was presented on the screen for three seconds, highlighting the current block’s target orientation and presence rate again. From the subject’s point of view, other than a contrast increase in the backward mask (titrated for each subject), everything remained the same between priming and main task trials. However, unbeknownst to the subjects, gratings were presented 50% of the time irrespective of the instructed target-presence rate (and contrary to the priming part of the block). Critically, the increased perceptual ambiguity due to strong backward masks allowed us to keep the grating presence rate the same between conservative and liberal priming blocks, while introducing a bias in the subject’s perceptual decisions.

**Figure 2.**
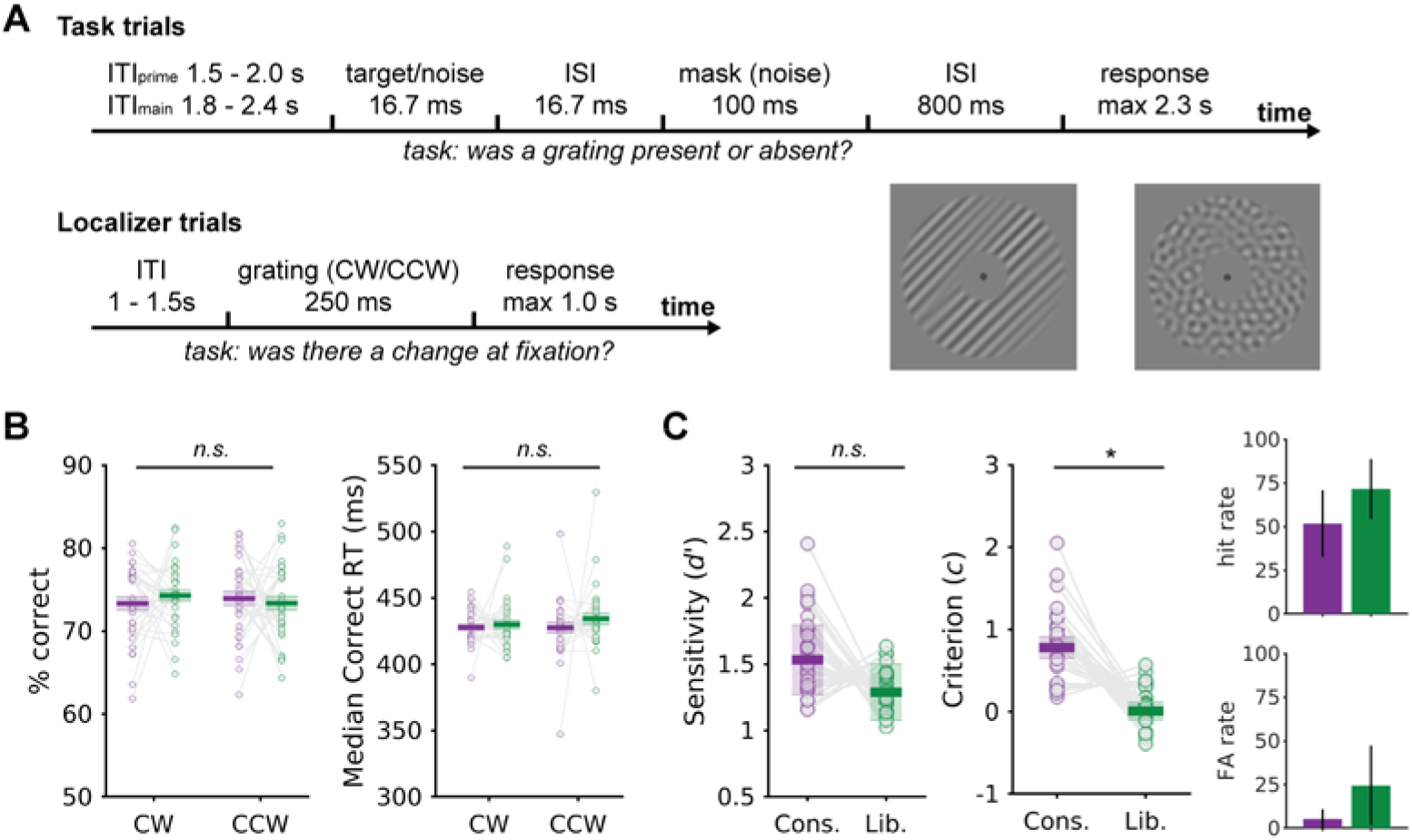
Experimental paradigm and behavioral results. **(A)** Schematic of the task trials (upper panel) and localizer trials (lower panel). Example grating and noise stimuli are shown for illustration purposes. **(B)** Accuracy and reaction times and **(C)** sensitivity and criterion for the main task trials during the MEG recording, and the corresponding hit and false alarm rates (mean and between-subjects standard deviations). Shaded areas denote within-subject standard errors in (B), and the variance of the group-level pooled *d’* and *c* in (C). Dots represent individual subjects (N=32). Asterisk denotes statistical significance (*p* < 0.05).

#### Grating localizer block

Interleaved with the main task blocks, subjects performed three blocks of a grating localizer task (see Figure 2A for trial schematic). Each block consisted of 120 trials, among which 18 trials contained a brief fixation blink (i.e., the central fixation dot turned from black to white for 50 ms). The subject’s task was to detect these occasional fixation blinks and to press a button as soon as possible when they saw them. Clockwise and counterclockwise gratings identical to those presented during the main task were presented in a pseudorandom order for 250 ms, followed by a uniformly jittered ITI of 1 to 1.5 s. We used a longer presentation duration (relative to that used in the main task trials) to increase the signal-to-noise ratio in the stimulus-driven response. These trials enabled us to identify brain regions most responsive to the bottom-up stimulation by grating stimuli. Trials containing fixation blinks—which served exclusively to maintain the subject’s attention on the fixation point—were excluded from later analyses.

### Stimuli

A bull’s eye (outer black ring = 0.5° × 0.5° degree of visual angle (dva), innermost black dot = 0.25° × 0.25° dva) was presented at the center of the screen throughout each block as the fixation point. Subjects were instructed to always maintain fixation, and not to blink during the presentation of the stimuli. A 2-s fixation window was presented before the start of the first priming and the first main task trial in each block. Each trial started with a brief presentation of the target (16.67 ms), within which either an oriented grating (Michelson contrast: 40%, spatial frequency: 1 cycle per °, orientation: 45° clockwise (CW) or counterclockwise (CCW) relative to vertical, randomized spatial phase) or a bandpass-filtered noise patch (Michelson contrast: 40%; spatial frequency: 1 cycle per °; randomly generated in each trial) was shown in an annulus (inner radius = 1.5°, outer radius = 7.5°, contrast of the stimuli decreased linearly to 0 over the outer and inner 0.5° radius of the annulus) around the central fixation (see Figure 2A for an example). After a brief (16.67 ms) ISI window, a backward mask consisting of a bandpass-filtered noise patch (Michelson contrast: titrated for each subject; spatial frequency: 1 cycle per °; randomly generated in each trial) was presented for 100 ms (see Figure 2A for an example). After another ISI window of 800 ms, the letters “P” (for grating present) and “A” (for grating absent) were presented on each side of the screen respectively, indicating the response mapping of the current trial, which changed pseudo-randomly and unpredictably across trials to avoid motor preparation confounds. Subjects had a maximum of two seconds to provide their response (i.e., grating present vs. absent). The inter-trial interval (ITI) started as soon as subjects committed a response. The ITI duration was jittered across trials between 1.8 and 2.4 seconds for the main task trials, and between 1.5 and 2.0 seconds for the priming trials. Note that we titrated the contrast of the backward mask via a 3-down-1-up staircase procedure at the beginning of each session, such that overall accuracy in a detection task with 50% target presentation rate was comparable across subjects and across sessions.

### Data acquisition

Stimuli were displayed on an LCD screen during the training session and on a semitranslucent screen (1920 × 1080 pixel resolution, 120-Hz refresh rate) back-projected by a PROpixx projector (VPixx Technologies) during MEG recordings. The experiment was programmed with Psychtoolbox (Brainard, 1997) in Matlab (The Mathworks, Inc.) and ran in a Linux environment. Brain activity was recorded using a 275-channel axial gradiometer MEG system (CTF MEG Systems, VSM MedTech Ltd) at 1200 Hz in a magnetically shielded room. Six permanently faulty channels were disabled during the recordings, leaving 269 recorded MEG channels. Three fiducial coils were placed at the subject’s nasion and both ear canals, to provide online monitoring of the subject’s head position (Stolk et al., 2013) and to serve as anatomical landmarks for offline co-registration with structural MRI scans. Eye position and pupil size were recorded using an infrared eye tracker (EyeLink, SR Research Ltd., Mississauga, Ontario, Canada) during the MEG recordings. Upon completion of the MEG session, the subject’s head shape and the location of the three fiducial coils were digitized using a Polhemus 3D tracking device (Polhemus, Colchester, Vermont, United States). T1-weighted MRI scans were acquired using a 3T MRI system (Siemens, Erlangen, Germany). Earplugs with a drop of vitamin E were placed at the subject’s ear canals during MRI acquisition, to facilitate co-registration with MEG data.

### Behavioral data analysis

We focused our analysis on main task trials (i.e., trials with high-contrast backward masks) recorded during the MEG session. Besides accuracy and reaction times (RT), our main behavioral variables of interest were sensitivity (*d*’) and criterion (*c*), which were estimated following Signal Detection Theory (SDT), assuming the internal signal and noise distributions shared the same variance (Macmillan & Creelman 2004) as follows:

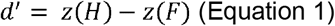

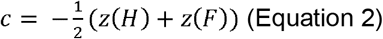

where *H* and *F* are the hit and false alarm rates, and *z*(*X*) denotes the inverse of the normal cumulative function evaluated at *X*. Moreover, the variances of these estimates are defined as:

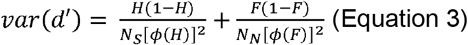

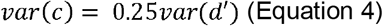

where *N_S_* and *N_N_* are the number of signal (S) and noise (N) trials, and *ϕ*(*X*) is the height of the normal density function at *z*(*X*).

To compare *d’* and *c* between priming conditions at the group-level, we first computed for each condition the *pooled d’* and *c* estimates as well as their corresponding variances, using the averaged hit rate (*H*) and false alarm rate (*F*) across subjects. This method results in more robust and unbiased group-level *d’* and *c* estimates compared to that estimated by averaging across subjects (Macmillan & Kaplan, 1985). We then calculated the difference in *pooled d’* (and *c*) between conditions as well as the confidence interval of the difference. If zero falls outside the 95% confidence interval of the *pooled d’* (or *c*) difference, then the conditions are considered significantly different from each other in *d’* (or *c*). Additionally, to assess the effects of priming and target orientation on accuracy and correct RTs, we used within-subject repeated-measures ANOVA (priming type: conservative vs. liberal; grating orientation: clockwise vs. counterclockwise).

To capitalize on trial-by-trial fluctuations and to capture the relationship between prestimulus state and behavioral outcomes, we adapted and extended the Generalized Linear Model (GLM) formulation of Signal Detection Theory (DeCarlo, 1998).

For each subject, we defined their responses (target presence/absence) as determined by a combination of factors (stimulus presence, the oscillatory state of interest, and the interaction between them) via a probit link function:

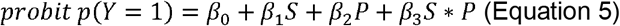

with *Y* the subject’s “target absence” and “target presence” responses (coded as zeros and ones), *S* the grating absence or presence (coded as zeros and ones), and *P* the trial-by-trial continuous measure of oscillatory power. Before feeding the oscillatory power of interest into the model, we first log-transformed it to reduce the skewness of the observed data, and then *Z*-transformed the corresponding output to ensure that the dynamic range is comparable across subjects. In this model, *Z*-transformed hit rate can be expressed as:

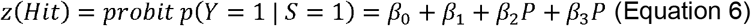

and the *Z*-transformed false alarm rate can be expressed as:

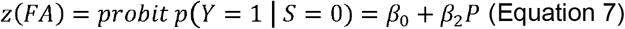

Replacing z(Hit) and z(FA) in Equation 1 with Equation 6 and 7 results in:

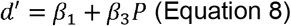

in which *β*_3_ describes how changes in oscillatory power of interest contribute to changes in sensitivity. Similarly, replacing z(Hit) and z(FA) in Equation 2 results in:

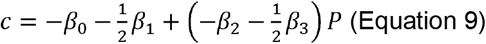

in which 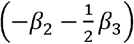 describes how changes in oscillatory power of interest contribute to changes in criterion. For convenience, we defined the effect of oscillatory power on sensitivity as

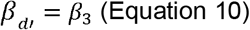

and the effect of oscillatory power on criterion as

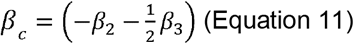

in later analysis and data visualization.

At the group-level, we extended the above model and constructed a generalized linear mixed model (GLMM) to better account for between-subject variation. Specifically, we fitted the model with “subjects” as the only random grouping factor, and included for each fixed effect its corresponding random slope coefficient and intercept. This was done using Matlab’s *fitglme* function with *quasinewton* optimizer. We reported results of the estimated coefficients *β_d’_* and *β_c_* and the corresponding statistical significance. Intuitively, a negative value of *β_d’_* (or *β_c_*) indicates that *d’* (or *c*) decreases as oscillatory power of interest increases. To obtain a null distribution of the *F*-values corresponding to *β_c_* and *β_d’_*, we fit the GLMM for all parcels, 100 times using shuffled data, in which the oscillatory alpha power time courses across trials were randomized and thus the temporal (autocorrelation) structure within a trial was preserved. We then used cluster-based permutation tests to evaluate the statistical significance of the fitted *β_c_* and *β_d’_*.

### MEG data analysis

#### MEG preprocessing

MEG data were preprocessed offline and analyzed using the FieldTrip toolbox (Oostenveld et al., 2011) and custom-built Matlab scripts. Trials of the main task blocks and grating localizer blocks were segmented and processed separately. All data were down-sampled to 400 Hz, after applying a notch filter to remove line noise and harmonics (at 50, 100, and 150 Hz). Trials with excessive noise were rejected via visual inspection before independent component analysis (ICA). ICA components were visually inspected and those representing eye and heart artifacts were then projected out of the data (Jung et al., 2000). Finally, outlier trials with extreme variance were removed, leaving on average 297 out of 320 (SD = 14.6) priming trials, 595 out of 640 (SD = 33.5) main task trials, and 331 out of 360 (SD = 18.0) localizer trials per subject.

#### MRI processing

MRI data were co-registered to the CTF coordinate system using the fiducial coils and the digitized scalp surface. Volume conduction models were constructed based on single-shell models of individual subjects’ anatomical MRIs (Nolte, 2003). Dipole positions were defined using a cortical surface-based mesh with 15784 vertices created using Freesurfer v6.0 (RRID: SCR_001847) and HCP workbench v1.3.2 (RRID: SCR_008750). The vertices were grouped into 374 parcels based on a refined version of the Conte69 atlas (Van Essen et al., 2012), allowing us to reduce the dimensionality of the data (similar to (Schoffelen et al., 2017). For each dipole position, lead fields were computed with a reduced rank, which accommodates the fact that MEG is blind to radial sources.

#### Event-related fields

Before calculating the event-related fields (ERFs), singe-trial data were baseline-corrected using a time window of [–0.5, 0] s for the priming and main task trials, and [–0.2, 0] s for the grating localizer trials. To avoid the confounding influence of noise (in the planar transformation) due to unequal trial numbers across conditions, trial numbers were equated via subsampling. Specifically, we subsampled an equal number of trials from each condition before averaging over trials, such that the number of trials per condition matched that in the condition of the fewest trials. Planar gradients of the MEG field distribution were then calculated, which makes spatial interpretation of the sensor-level data easier and facilitates comparison of ERF topographies across subjects. We repeated the above-mentioned procedure 10 times per condition, to ensure every trial was used at least once. We then averaged across all corresponding planar-combined averages to obtain ERFs per condition.

#### Spectral Analysis

Time-frequency representations (TFRs) of power were calculated for each trial by applying a fast Fourier transform to short sliding time windows. We applied Hanning tapers of 4-cycles length in time steps of 50 ms to single-trial data, prior to computing the spectra (4–30 Hz). For sensor-level analyses, spectral decomposition was applied to synthetic planar gradient data, and combined into single spectra per sensor. For visualization purposes, condition-average TFRs were expressed relative to a frequency-specific baseline window, which started 4 cycles and ended 2 cycles before stimulus onset (e.g., for 10-Hz activity, a 400 to 200 ms prestimulus time window was used as baseline window) to prevent leakage of post-stimulus activity into the baseline window. Note that for all statistical analyses, non-corrected TFR data were used. To compute TFRs of an anatomical parcel (source-level), spectral decomposition was performed on the parcel’s response time series (see *Source reconstruction*). To account for intra- and interindividual variability in alpha peak frequency (Haegens et al., 2014), we defined the individual alpha peak frequency for each brain area (parcel) and each subject separately. This was done with the priming trials by computing the Fourier spectrum of each parcel’s pre-stimulus 1-s time series and finding the local maximum within the 7–14 Hz band of the resulting Fourier spectrum. We computed the oscillatory alpha power within each parcel by averaging the above-mentioned power estimate within a ±1 Hz band centered at the individual alpha peak frequency. We checked the prevalence of alpha oscillatory activity by counting the numbers of subjects exhibiting prominent alpha oscillatory activity identified using the above-mentioned method (Figure 3A). To estimate the time courses of oscillatory alpha phase, we used Hanning tapers of 200 ms length (zero-pad to one second) at the individual alpha peak frequency, to slide over each anatomical parcel’s response time series in steps of 5 ms. The phase was computed as the angle of the resulting Fourier coefficients. Unless otherwise specified, analyses linking alpha oscillatory power/phase to behavior are all based on that of individualized alpha peak frequencies.

**Figure 3.**
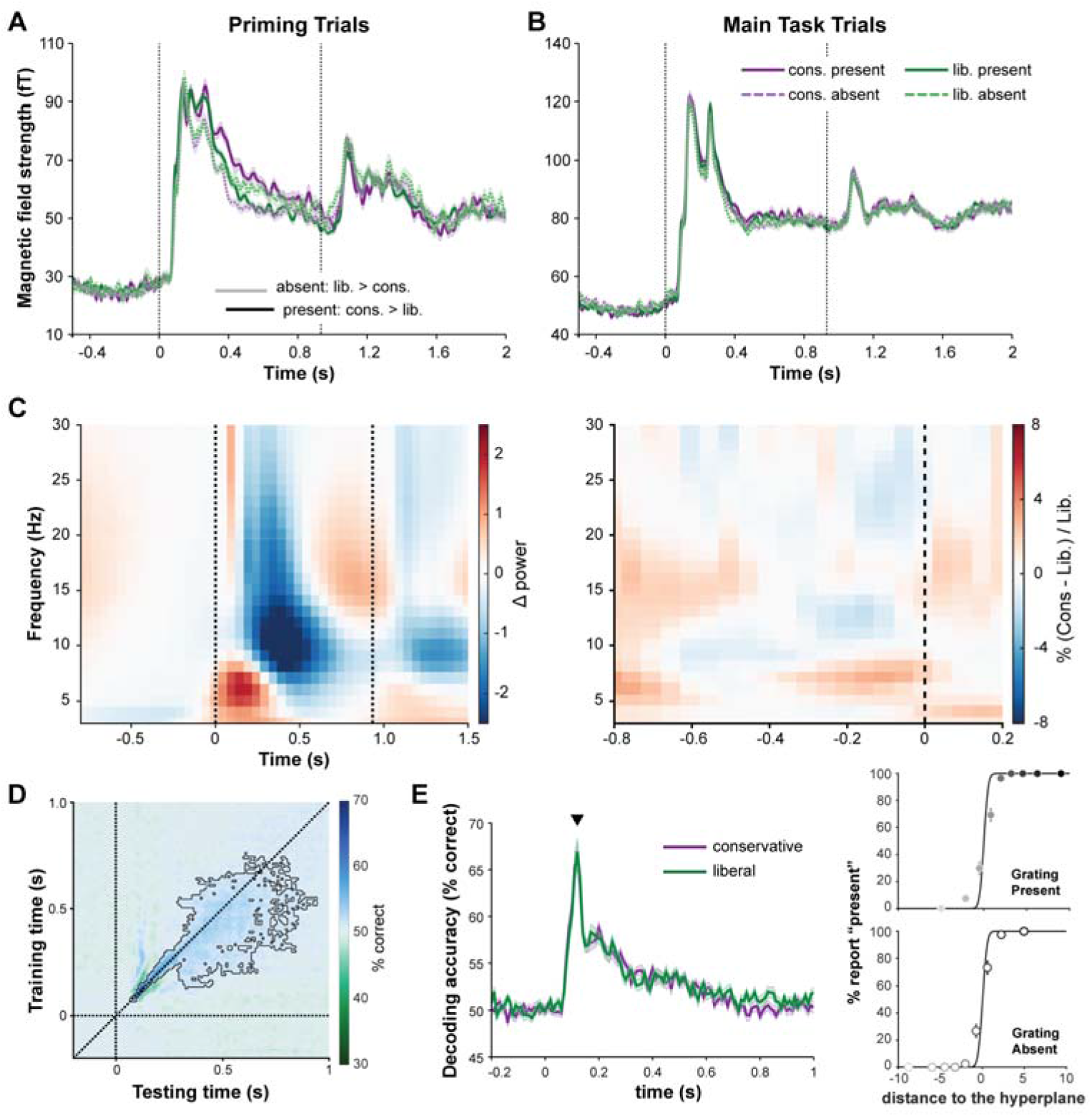
Event-related fields (ERF) and time-frequency representation (TFR) of selected sensors and classification results. **(A)** ERFs for the priming trials (showing data for of the 20 selected sensors most responsive to the localizer gratings). Black and grey lines at the bottom indicate statistically significant time points identified by cluster-based permutation tests. **(B)** Same as (A) for main task trials. **(C)** TFR of the main task trials (left panel) and the TFR of the difference between conservative and liberal trials (right). **(D)** Temporal generalization matrix of classification accuracy. **(E)** Time courses showing the diagonal of the temporal generalization matrices. The right panels show subjects’ response to grating-present and absent trials as a function of the amount of evidence in the classifier trained and tested at time point 0.12 s (marked by a triangle above the time series). Dots represent group averages of each bin.

#### Phasic modulation of accuracy

We assessed whether behavioral accuracy, both that of the subjects and the neural decoder (see *Multivariate pattern analysis*), changed as a function of oscillatory alpha phase. We grouped the trials based on their corresponding phase estimates of a specific time window and parcel into 16 equidistant phase bins (bin centers = [0, 22.5,…, 337.5] degree, bin width = 45 degree), and computed the average accuracy for each bin. We combined all trials (conservative and liberal) in this analysis because it is unlikely that subjects endogenously aligned their instantaneous alpha phase with respect to the stimulus onset, as onset was randomly jittered and hence unpredictable to the subjects. We used accuracy as an index of interest for the current analysis because accuracy, compared to *d’*, is a more robust measure of performance when trial number is limited (in this case, less than 80 trials per phase bin). We then fitted a cosine function of one cycle and unknown amplitude and phase to the resulting 16 accuracy scores. The amount of phasic modulation of accuracy was defined as the fitted amplitude of the cosine function. Intuitively, an estimated amplitude of 3% suggested that the model predicted an accuracy difference of 6% between trials of the optimal phase and those of the suboptimal phase. This procedure was applied to every time window and parcel of interest. To obtain a null distribution of the fitted amplitudes for statistical inference, we circularly shifted the phase time course of every trial with a random value between −360 to 360 degrees before phase binning the data, which allows randomization without breaking the temporal (autocorrelation) structure within a trial.

#### Source reconstruction

Source-level analyses reported in the results section are based on a sample of 29 subjects, whose T1-weighted anatomical images were available. The linearly constrained maximum variance beamformer approach (Van Veen et al., 1997) was used to obtain the source reconstruction of the event-related response to the localizer gratings. The data covariance matrix was computed over a window of [−1.0, 1.0] s for priming and main trials and [−0.5, 0.5] s for localizer trials, time-locked to stimulus onset, and was subsequently used to construct common spatial filters. To estimate the amplitude of the single-trial stimulus evoked response, we projected the trial data computed over [0, 0.2] s through the spatial filters, and then normalized the resulting amplitude by that of a baseline window of [−0.2, 0] s time-locked to stimulus onset. Neural response time series of an anatomical parcel were computed by taking the first principal component of the time series over all dipole positions within the parcel.

We used the partial canonical coherence beamformer approach (Schoffelen et al., 2008) to localize the sources of oscillatory alpha-band activity in response to the main task trials. To estimate the spatial distribution of alpha-band power, we first extracted 500-ms data segments (time window = [−0.5 0], time locked to stimulus onset), then computed cross-spectral density (CSD) matrices using Slepian tapers (Mitra & Pesaran, 1999) centered at a frequency of 10 (±4) Hz. With the CSD matrices and the lead fields, a common spatial filter (for all trials across both conditions) was constructed for each dipole position for each subject. By projecting sensor-level CSD through the common spatial filter, the spatial distribution of power was then estimated for each trial. To reduce data dimensionality, we took the mean estimated power of all dipoles within each parcel as the single-trial power estimate of the parcel.

#### Multivariate pattern analysis

To probe the representational content of neural activity, we performed multivariate pattern analysis using the MVPA-light toolbox (Treder, 2020). Before running the classification analysis, we first applied a low-pass filter at 30 Hz and down-sampled the data to 100 Hz. Multivariate logistic regression (L2 regularized) was then applied to predict the grating’s presence (present vs. absent), based on the sensor-level spatial distribution of neural activity at each time point. To avoid overfitting, we trained the classifier with priming trials, and tested on the main task trials. This procedure was done within-subject in a time-resolved manner, resulting in a temporal generalization matrix (King & Dehaene, 2014).

#### Cluster-based permutation tests

Statistical significance was evaluated using cluster-based permutation tests (Maris & Oostenveld, 2007) at the sensor-level. The time interval of interest was [−0.8, 0] (i.e., the 800-ms window before stimulus onset). For ERFs, data of different preselected sensors were first combined, resulting in a single time series per subject. These time series data were then compared univariately at each time point. Neighboring time points for which a two-tailed paired *t*-test (or a repeated-measures ANOVA) resulted in a nominal *p*-value smaller than 0.05 (uncorrected) were clustered. A similar procedure was applied to the time-frequency representations (TFRs), with clustering across spectral, temporal and spatial dimensions. The sum of the *T*-values (or *F*-values) within a cluster was then computed as cluster-level statistics. The cluster with the maximum sum was subsequently used as test statistic. By randomly permuting the condition labels (or by shuffling the predictor variable of interest across trials) and recalculating the test statistic 10,000 times (or 100 times, for computationally expensive operations such as the GLMM and phasic modulation analysis), we obtained a reference distribution of maximum cluster *T*-values (or *F*-values) to evaluate the statistic of the actual data (alpha = 0.05). For reference, we also report the smallest *p*-value obtained for cluster-based permutation tests that failed to reject the null hypothesis.

## RESULTS

### Priming leads to criterion shifts

Subjects finished eight blocks of the main experimental task and three blocks of a localizer task during the MEG session. For the localizer task, they had to detect and report as fast as possible occasional fixation changes, which occurred in 15% of all trials. Subjects correctly reported fixation changes on 99.5% (between-subjects SD = 0.5 %) of all trials with mean median reaction time (RT) of 406 ms (between-subjects SD = 49 ms), confirming close engagement in the task.

For the main experimental blocks, subjects’ task was to report the presence (or absence) of an oriented grating as accurately as possible. They consistently performed at ceiling during the priming phase when the masking effect was minimal (mean accuracy = 95.2%, with a between-subjects SD of 5.1%; mean median RT = 421 ms, with a between-subjects SD of 73 ms). For the main task trials, in which the mask was presented at a contrast level titrated for each subject, overall accuracy dropped to around 75% as aimed for. Note the mean accuracy was lower than that expected from a 3-down-1-up staircase procedure (i.e., about 80%), likely because the instructed target-presence rate was different from the actual target-presence rate. Two-way (priming type × grating orientation) repeated-measures ANOVA showed no significant differences across conditions in RT or accuracy (*F*s < 1; Figure 2B). Results of the SDT analysis (Figure 2C) showed that subjects were significantly more liberal in reporting “target present” in the liberal priming blocks than in the conservative priming blocks (*p* < 10^−6^), validating our experimental manipulation. Critically, their sensitivity (*d’*) was comparable between the liberal and conservative blocks (*p* = 0.082). Taken together, our behavioral results show that the current paradigm successfully introduces criterion shifts in subjects’ detection responses.

### Priming does not lead to modulation of pre-stimulus oscillatory power or stimulus-evoked activity

According to the additive hypothesis, pre-stimulus alpha power modulates the decision criterion. Therefore, one may expect (top-down) changes in criterion induced by different priming conditions to be reflected in alpha power differences. We first compared pre-stimulus timefrequency representations (TFRs) between the conservative vs. liberal trials, without making any assumptions about the frequency range of potential effects. We observed no significant differences in pre-stimulus TFRs (*p* = 0.366, cluster-based permutation test with frequencies from 1 to 30 Hz, time points from 800 ms pre-stimulus to stimulus onset, including all sensors) between the two priming conditions. We then repeated our statistical tests by constraining our frequencies of interest to around the alpha range (6 – 14 Hz), or by constraining the channels of interest to 20 subject-specific channels with strongest evoked response to the localizer gratings (in a time window of [0.1, 0.2], relative to baseline). We found no significant pre-stimulus TFR differences between the two priming conditions in either analysis (Figure 3C).

In addition to pre-stimulus activity, we also examined whether priming modulates the sensory response amplitude to the stimuli. We constrained these analyses to the abovementioned 20 subject-specific channels, which primarily reflect the stimulus-evoked responses. We did not observe any significant differences in the average event-related fields (ERF) amplitude (averaged across all selected sensors within each subject) between different priming conditions for the main task trials (Figure 3B). Interestingly, when the same contrasts were repeated for the priming trials, we found that for target-present trials, conservative priming led to larger ERF amplitude 400 to 650 ms after stimulus onset (*p* < 10^−4^; Figure 3A solid lines) compared to liberal priming; and this pattern was reversed for target-absent trials (*p* < 10^−4^; Figure 3A dotted lines). These observations may reflect a general P3 surprise effect (Mars et al., 2008) to the less frequently presented stimuli, given that target-present and target-absent trials were rare events (i.e., 20% of trials) in the conservative and liberal priming blocks respectively. These ERF results also suggest that subjects incorporated the different probabilistic information in sensory processing during the priming phase.

Taken together, we show for the main task trials that neither the pre-stimulus oscillatory activity nor the stimulus-evoked activity reflects the (top-down) criterion changes introduced by priming.

### Pre-stimulus alpha power modulates *d’*: region of interest analysis

If excitability modulation by alpha oscillatory power has any behavioral consequence, its magnitude immediately prior to stimulus onset, within the neuronal populations most responsive to the stimuli, should be most relevant for subsequent detection responses. Following this rationale, we zoomed in on the anatomical parcel most responsive to localizer gratings (defined for each subject, see Figure 4B for spatial topography of the selected parcels), and asked whether and how alpha power immediately before the stimulus onset in these functionally defined visual regions of interest (ROIs) predicts the subject’s response. While we observed prominent oscillatory activity within the alpha-band during the pre-stimulus time window (Figure 4B, right panel), we found no significant difference between the two conditions (i.e., conservative vs. liberal) in the pre-stimulus TFR (cluster-based permutation test with frequencies from 1 to 30 Hz, time points from 800 ms pre-stimulus to stimulus onset: *p* = 0.257) nor in the power spectrum (cluster-based permutation test with all frequencies: *p* = 0.078).

**Figure 4.**
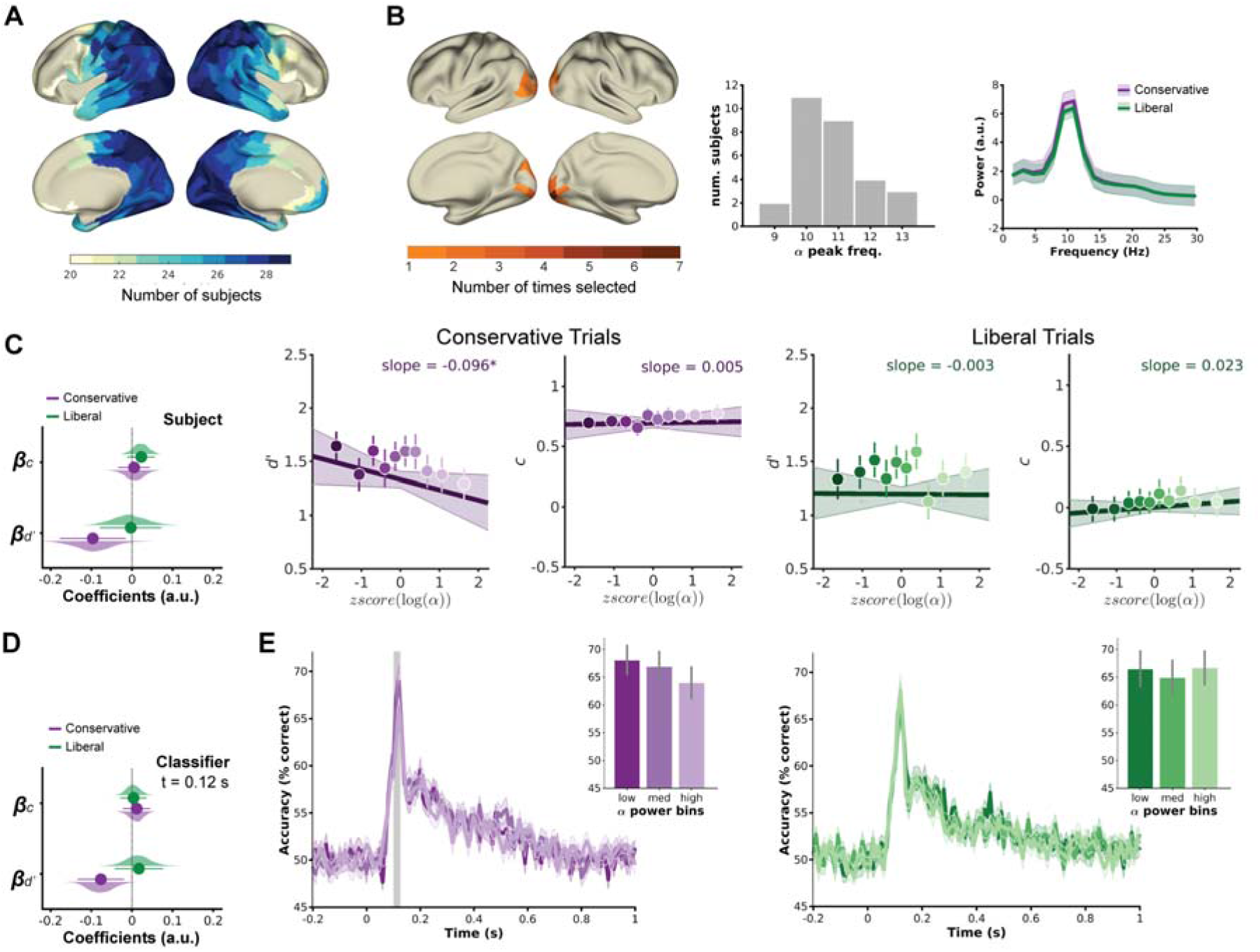
ROI-based analysis of relationship between alpha and performance. **(A)** The prevalence (indexed by the number of subjects) of alpha oscillatory activity across brain regions. **(B)** Spatial topography of the visual regions of interest most responsive to grating stimuli, and the corresponding alpha peak frequencies and the pre-stimulus power spectrum. **(C)** Left: GLMM results: estimated coefficients and the corresponding 95% confidence intervals, and the distribution of bootstrap estimates for the parameters. Middle: visualization of the fitted coefficients, for the conservative priming trials. Dots represent group-level average estimates and the corresponding within-subject standard errors. The line shows the fitted model. Right: same as the middle panel, but for the liberal trials. **(D)** GLMM results: estimated coefficients and the corresponding 95% confidence intervals, and the distribution of bootstrap estimates for the parameters. **(E)** Decoding accuracy (diagonal of the temporal generalization matrix) as a function of pre-stimulus alpha power for conservative (left) and liberal (right) trials. Gray shaded area marks the time window during which pre-stimulus alpha power significantly modulated decoding accuracy indicated by the corresponding cluster-based permutation test. Shaded area around the time series denotes within-subject standard error. Bar graphs show mean decoding accuracy (within the gray shaded area) as a function of pre-stimulus alpha power. Error bars denote within-subject standard error.

Next, we asked: do trial-by-trial changes of pre-stimulus oscillatory alpha power influence the subject’s detection responses? We applied generalized linear mixed models (GLMMs; see Methods) which account for both between-subjects and within-subject trial-by-trial response variability to address this question. Our modeling results (Figure 4C) showed a significant negative effect of pre-stimulus alpha power on *d’*, which was exclusively driven by conservative trials (conservative: *β_d’_* = −0.096 (bootstrap SD = 0.030), *p* = 0.021; liberal *β_d’_* = −0.003 (bootstrap SD = 0.031), *p* = 0.939). Specifically, subjects’ *d’* decreased as alpha power increased under the conservative priming condition. The effects of pre-stimulus alpha power on criterion were not statistically significant (conservative: *β_c_* = 0.005 (bootstrap SD = 0.017), *p* = 0.798; liberal *β_c_* = 0.023 (bootstrap SD = 0.017), *p* = 0.197). Collectively, our results indicate that pre-stimulus alpha power in visual regions most responsive to the target stimuli modulated perceptual sensitivity, not criterion.

### Pre-stimulus alpha power enhances sensory representation

To evaluate the quality of visual information coding, we used multivariate pattern analysis (MVPA), operationalizing the quality of visual representation as the neural classifier’s classification performance. We first confirmed that classifiers trained on priming trials (classifying target-present vs. target-absent trials) are generalizable (i.e., performed significantly above chance) to the main task trials (Figure 3D). Zooming in to the diagonal of the matrix, we found that classification accuracy did not differ significantly between priming type (*p* = 0.092, cluster-based permutation test on time window [0, 1.0] s, Figure 3E). Moreover, we found the subjects’ responses can be predicted by the classifier’s decision evidence (i.e., distance to the hyperplane) at the time classification accuracy peaked (Figure 3E). Hence, the neural classifiers trained on priming trials not only generalizes to main task trials but also tracks subjects’ responses quantitatively and qualitatively.

We examined whether pre-stimulus alpha power modulates the quality of visual information coding by first focusing on the neural classifier’s response at the time when classification accuracy peaked (trained and tested at the time point 120 ms after stimulus onset). Similar to the above-mentioned analyses linking pre-stimulus alpha power with the subject’s behavior, we applied the GLMM to the classifier’s responses. We found a negative correlation between pre-stimulus alpha power and the classifier’s *d’* for the conservative trials (Figure 4D, conservative: *β_d’_* = −0.077 (bootstrap SD = 0.029), *p* = 0.009, *β_c_* = 0.012 (bootstrap SD = 0.017), *p* = 0.495; liberal: *β_d’_* = 0.017 (bootstrap SD = 0.031), *p* = 0.587, *β_c_* = 0.004 (bootstrap SD = 0.017), *p* = 0.823), suggesting that under the conservative priming condition lower pre-stimulus alpha power leads to enhanced task-relevant information coding.

We next asked whether the modulatory effect on classification accuracy by pre-stimulus alpha power manifested at time points other than when classification accuracy peaked. For each priming condition, we divided trials into low, medium, and high alpha power bins (with about 100 trials per bin, allowing for robust estimates of classification performance), using the pre-stimulus alpha power within our visual ROIs. We then compared the classification accuracy time course across the three bins using cluster-based permutation test (on the diagonal of the temporal generalization matrix, time window = [0, 0.5] s). This analysis showed that lower pre-stimulus alpha power in the visual ROIs was associated with higher classification accuracy about 120 ms after target onset, and that this modulation was present only for the conservative trials (*p* = 0.038, Figure 4E), not the liberal ones (*p* = 0.159).

Together, our classification results show that a state of decreased pre-stimulus alpha power leads to enhanced visual representation, which likely results in better detection sensitivity.

### Pre-stimulus alpha power modulates *d’* predominantly in occipital-parietal areas: wholebrain analysis

While the above ROI analysis enabled us to maximize statistical power to address our research questions, it may have precluded us from observing interesting effects taking place outside our visual ROIs. To explore whether and how pre-stimulus alpha power fluctuations in different brain regions modulated subjects’ detection responses, we applied the GLMM analysis to all anatomical parcels, and asked whether alpha power changes in any of them modulated the subject’s *d’* or criterion. We applied cluster-based permutation tests to correct for multiple comparisons across time (from 600 to 200 ms pre-stimulus) and space (374 anatomically-defined parcels, whole brain). Under the conservative priming condition, five parcels along the left ventral visual pathway showed significant negative alpha power modulation on *d’* (*p* < 0.01; Figure 5A), whose effect was most pronounced at time points immediately before stimulus onset. Under the liberal priming condition, a cluster of about 20 parcels extending bilaterally from the somatosensory cortex to superior parietal lobule showed significant negative alpha power modulation on *d’* (*p* < 0.01; Figure 5B). In contrast, we observed a positive criterion modulation by alpha power for brain regions overlapping with this cluster, now for the conservative priming condition (*p* < 0.01; Figure 5C); though this modulation disappeared for time points immediately preceding the stimulus presentation. There was no significant criterion modulation by alpha power under the liberal priming condition.

**Figure 5.**
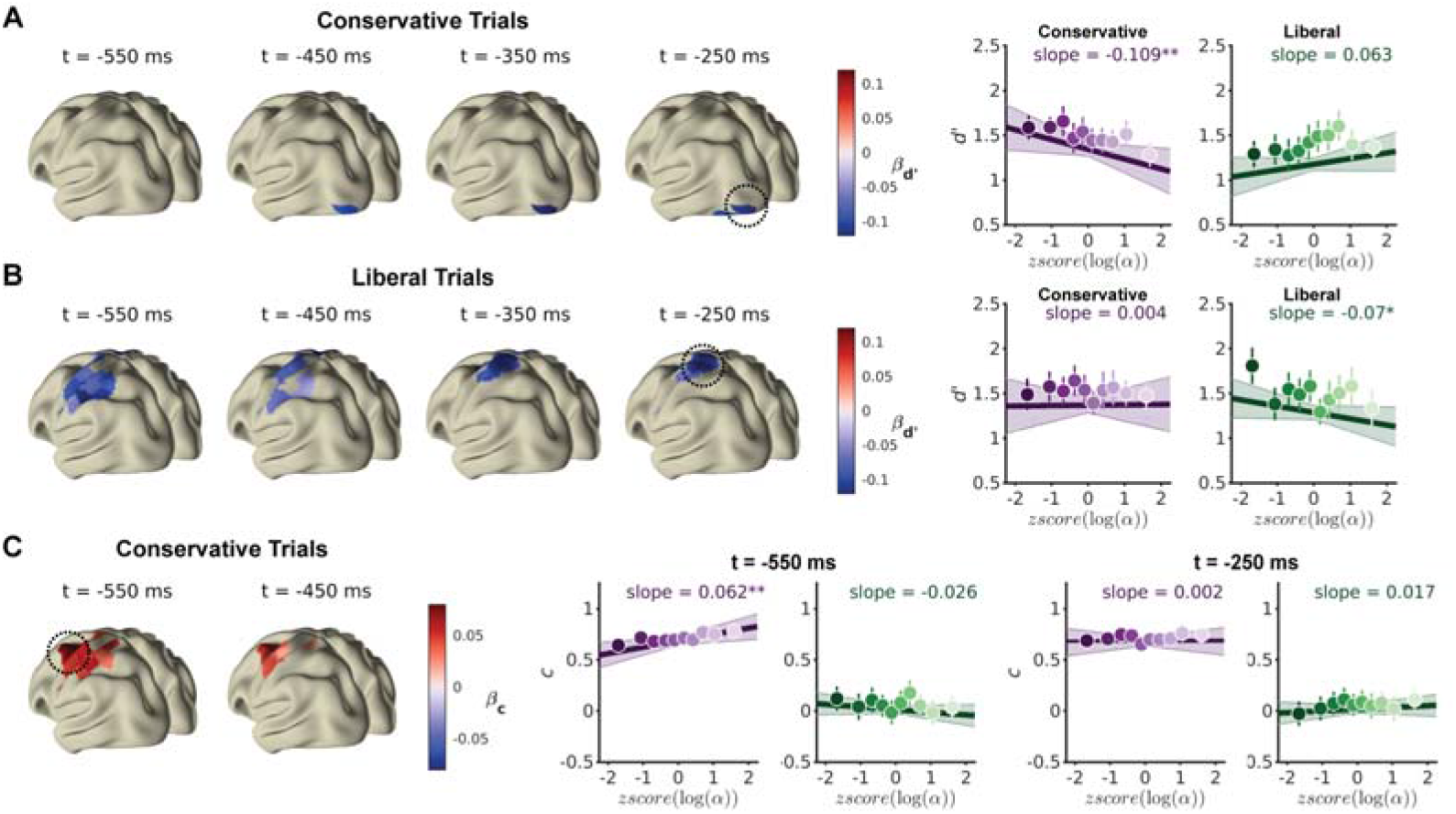
Whole-brain GLMM analysis of relationship between alpha and performance. **(A)** Estimated *β_d’_* for the conservative trials (masked by statistical significance after the correction of multiple comparisons) across pre-stimulus time points, and the zoomed-in visualization of the fitted coefficients of the highlighted parcel at t = −250 ms. **(B)** Similar to (A), but for the liberal trials. **(C)** Similar to (A), but for estimated *β_c_* for the conservative trials. The highlighted parcel’s fitted coefficients at both t = −550 ms and t = −250 ms are shown.

### Pre-stimulus alpha phase modulates accuracy in a phasic manner

Finally, we asked how detection accuracy fluctuated as a function of oscillatory alpha phase. We estimated the extent of phasic modulation of accuracy by taking the amplitude of the fitted cosine function that best described accuracy changes as a function of oscillatory alpha phase. We first examined the fitted cosine’s amplitude over time during the pre-stimulus time window in the functionally-defined visual ROIs. The estimated amplitudes were significantly above chance around 200 ms before the stimulus onset, suggesting significant phasic modulation of detection accuracy (Figure 6AB). Interestingly, such a phasic modulation of the subject’s accuracy seemingly faded out immediately before the stimulus onset. In contrast, the phasic modulation of the neural classifier’s responses peaked at around 100 ms immediately before stimulus onset, suggesting that oscillatory alpha phase in these visual ROIs may have different impacts on the sensory representation and the subject’s detection responses.

**Figure 6.**
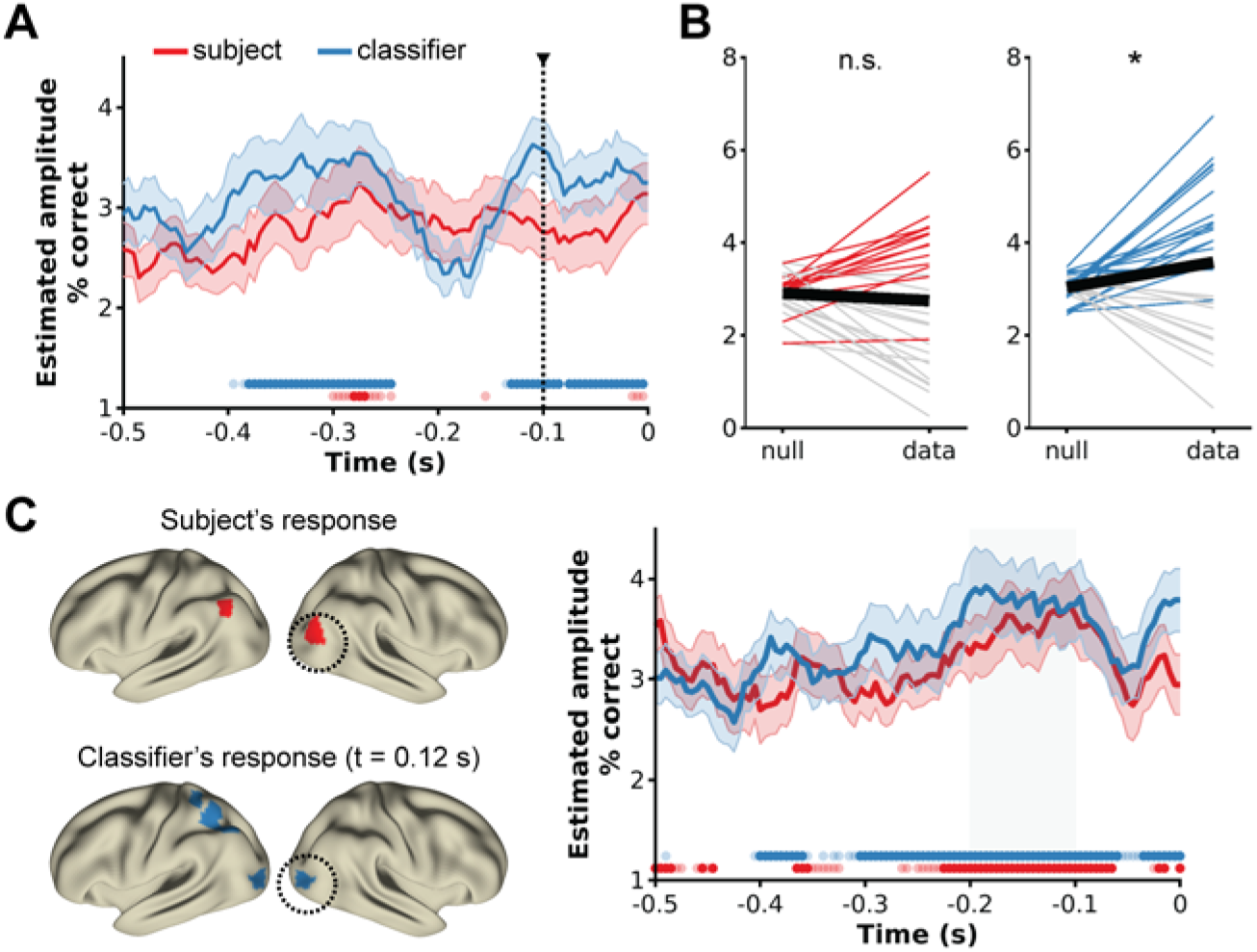
Phasic modulation of accuracy. **(A)** Estimated amplitude of the phasic modulation of subjects’ and neural classifier’s accuracy over time. Shaded areas around the curve denote between-subjects standard deviations, and dots at the bottom denote statistical significance, of which the lighter ones correspond to *p* < 0.05 uncorrected, and the darker ones correspond to those corrected (FDR = 0.05) for multiple comparisons. **(B)** Estimated amplitude of the phasic modulation of subjects’ and neural classifier’s accuracy at the time point 100 ms before stimulus onset in the functionally-defined ROIs. Thinner lines denote data from individual subjects, with the gray ones denoting cases where the estimated amplitude failed to exceed that of the null distribution. Thick black lines denote the group average amplitudes. **(C)** Left: Parcels showing significant phasic modulation on the subjects’ accuracy and classifier’s accuracy (when trained and tested at 0.12 s post-stimulus onset). Right: of the highlighted (circled) parcels in the left panel, the corresponding time courses of the estimated amplitude. Same conventions as in (A). Grey shaded area denotes the [−0.2, −0.1] time window used by the left spatial topography.

Next, we asked whether any anatomical parcels exhibited significant phasic accuracy modulation by oscillatory alpha-band activity. We focused on the fitted cosine’s amplitude corresponding to the time window immediately before stimulus onset whose phase estimates were unaffected by post-stimulus responses ([−0.2, −0.1] s). After correcting for multiple comparisons (false discovery rate 5%), two occipital parcels in the dorsal visual stream displayed significant phasic modulation of the subject’s detection response, while several parcels along the dorsal visual stream extending towards the superior parietal lobe (Figure 6C) displayed significant phasic modulation of the classifier’s peak response (i.e., classifier trained and tested at 120 ms post-stimulus).

These analyses of oscillatory alpha phase suggest that accuracy was modulated in a phasic manner, and that such phasic modulations are most likely driven by occipital-parietal alpha phase along the dorsal visual pathway.

## DISCUSSION

To understand how alpha oscillations modulate perception, we recorded MEG in human subjects while they performed a visual detection task. We found that criterion shifts induced by our priming manipulation were not reflected in a modulation of pre-stimulus alpha activity. Rather, we found that trial-by-trial changes of pre-stimulus alpha power modulate perceptual sensitivity, and that alpha phase in regions along the dorsal visual pathway modulates behavioral accuracy.

By providing the target-presence rate and presenting clearly visible targets at the start of each block (i.e., the priming trials), we successfully changed subjects’ perceptual expectations and thereby introduced criterion shifts in their decisions. While recent work has demonstrated that the human brain implements perceptual expectations by potentiating neural activity representing the expected stimuli (Ekman et al., 2017; Kok et al., 2017), we did not observe such a potentiation, nor did we observe any neural activity difference between the two priming conditions. Unlike a previous study where criterion was manipulated via changing the stimulus-response reward contingencies (Kloosterman et al., 2019), our current paradigm minimized the effect of reward contingency on neural activity and arguably allowed a cleaner reconstruction of sensory responses. Moreover, we manipulated subjects’ perceptual expectations in a block-wise manner, therefore our null finding cannot be explained by the brain not having enough time to flexibly adjust its internal state. As indicated by the significant and sustained priming effect on behavior, subjects incorporated different perceptual expectations consistently throughout the recording session. However, the endogenous shift of decision criterion based on different perceptual expectations does not seem to require or rely on pre-stimulus alpha modulations.

Next, we asked whether trial-by-trial fluctuations of pre-stimulus alpha oscillations lead to changes in criterion or sensitivity. By focusing first on the visual regions most responsive to our grating stimuli, we showed that both the subject’s perceptual sensitivity and information content decodable from the neural activity patterns are enhanced when pre-stimulus alpha power in early visual areas is low. Interestingly, the time window during which information content was modulated by pre-stimulus alpha power (i.e., about 120–160 ms post stimulus onset) overlaps with that during which local (as opposed to top-down) recurrent processes dominate (Wyatte et al., 2014), suggesting that lower pre-stimulus alpha power may lead to enhanced local recurrent processing. In line with a large body of spatial attention literature (e.g., Haegens et al., 2011; Thut et al., 2006; van Ede et al., 2011) showing that the extent of cue-induced alpha power lateralization correlates with behavioral performance, our current results extend previous work and reveal that ongoing trial-by-trial alpha power fluctuations predict sensitivity changes. Collectively, these findings support the multiplicative hypothesis, namely, that lower pre-stimulus alpha power amplifies the neural response such that the signal of interest becomes more distinct from noise, thereby improving perceptual sensitivity.

We also asked whether and how alpha oscillations in different brain areas may have different modulatory effects on behavior. We showed that trial-by-trial changes of alpha power within the ventral visual pathway predicted behavioral sensitivity when subjects were primed to adopt a more conservative detection criterion, and that fluctuations within the parietal and somatosensory regions predicted behavioral sensitivity when subjects were primed to adopt a more liberal criterion. These different spatial distributions may reflect that when subjects adopt different detection criteria, the processing bottleneck lies at different processing stages. When sensory information travels downstream, task-irrelevant information (i.e., noise) accumulates and scales to a larger extent compared to task-relevant information (i.e., signal), resulting in decreased signal-to-noise ratio of task-relevant information. If gating of sensory noise is ineffective at the start of sensory processing (e.g., in V1), the internal response used for perceptual decision-making is unlikely to reach the criterion later on, regardless of the effectiveness of gating of downstream noise. Therefore, when subjects adopt a more conservative criterion, alpha power fluctuations indexing the extent of effective gating in early visual areas become more determinant of perceptual sensitivity. Conceivably, while effective gating of upstream and downstream noise indexed by spontaneous alpha power fluctuations jointly determine perceptual decisions, it is possible that their respective importance is dependent on subjects’ decision strategy (i.e., criterion). This proposal is speculative, however, and in need of further empirical evidence.

Our whole-brain exploratory analysis further revealed that, under the conservative priming condition, trial-by-trial changes of pre-stimulus alpha power within the somatosensory regions predicted the subject’s detection criterion. Curiously, this relationship was most pronounced for alpha activity 500 ms preceding the stimulus onset, and disappeared for time points closer to the stimulus onset. The current observation appears to mirror previous reports showing that subjects adopt a more conservative criterion when alpha power increases (Iemi et al., 2017; Limbach & Corballis, 2016; Samaha et al., 2017), and it further suggests that prestimulus alpha power modulation on *d’* and criterion have different time courses. The current findings also highlight a methodological challenge faced by studies using conventional detection paradigms with variable pre-stimulus windows (Iemi et al., 2017; Limbach & Corballis, 2016), in that the definition of the “pre-stimulus” time window in such designs is arbitrary, as no stimulus is presented on target-absent trials and consequently no stimulus onset to reference to. Arguably, this can be reconciled if the stimulus onset is made predictable to the subject (e.g., by using temporal cues and fixed pre-stimulus windows). However, the immediate question is to what extent top-down processes contribute to pre-stimulus alpha fluctuations recorded using paradigms with predictable stimulus onsets. Such paradigms probably capture different neural dynamics than ours (i.e., ongoing fluctuations vs. top-down modulation), as previous work found that human subjects endogenously adjust their oscillatory alpha state (both in terms of power (Rohenkohl & Nobre, 2011) and phase (Samaha et al., 2015)) prior to the predictable onset of relevant stimuli to optimize sensory processing.

Finally, having demonstrated the relationship between alpha power and perception, we asked whether pre-stimulus alpha activity modulates behavioral accuracy in a phasic manner, as predicted by the inhibition hypothesis (Jensen & Mazaheri, 2010; Klimesch et al., 2007). We found that the alpha phase along the dorsal visual pathway immediately prior to stimulus onset modulated the subject’s and the classifier’s response accuracy. While one may expect the alpha phase effect to be dependent on and additive to the alpha power effect (Mathewson et al., 2009), our findings are consistent with previous work showing that alpha phase and power effects have different spatial topographies (Busch & VanRullen, 2010). Further research is needed to understand how the interplay between oscillatory alpha power and phase modulates perception.

To summarize, the current study shows with complementary univariate and multivariate results that changes of pre-stimulus alpha-band power modulate the quality of visual information coding at the neural level and perceptual sensitivity at the behavioral level. These findings provide important insights into how ongoing neural activity sets the internal state and subsequently influences perception.

## Acknowledgments

We thank Kristina Baumgart and Felix Linsen for their assistance with data acquisition. This study was supported by the Netherlands Organization for Scientific Research Vidi grants (NWO 016.Vidi.185.137 awarded to SH, and 864.14.011 to JMS), the Chinese Scholarship Council (CSC20170800036 awarded to YJZ), and the European Union Horizon 2020 Program (ERC Starting Grant 678286, “Contextvision” awarded to FPdL).

